# Nanopore sequencing identifies a distinct Tyr::CreER^T2^ allele circulating in existing mouse stocks

**DOI:** 10.64898/2026.07.29.741227

**Authors:** Leah Pomfret, Aro Nugawela, Emma L. Wilkinson, Barbara Shih, Richard L. Mort

**Affiliations:** Division of Biomedical and Life Sciences, Faculty of Health and Medicine, Lancaster University, Lancaster, UK

**Keywords:** Tyrosinase, melanocyte, melanoblast, melanoma

## Abstract

The Tyr::CreER^T2^ transgenic mouse lines are essential tools for conditional gene manipulation in melanocytes and are widely used in melanoma research. Two independent lines are in common use, the Bosenberg line and the Larue line, with transgene integration sites for both characterised by whole genome sequencing (Aktary et al., 2020). Precise knowledge of these sites is critical for designing complex genetic crosses. Here we report that a Tyr::CreER^T2^ mouse stock routinely used in *Braf*^*CA*^*;Pten*^*flox*^ melanoma models, and associated with elevated spontaneous melanoma penetrance, carries a previously undescribed integration on Chromosome 1, distinct from the previously reported Chromosome 2 integration. Using long-read Nanopore whole genome sequencing, we mapped the integration to an intergenic locus between *Alppl2* and *Alpi*, revealing a 2,431 bp genomic deletion at the insertion site. We developed position-specific junction PCR and qPCR assays to genotype this allele and confirmed that offspring are born at Mendelian ratios. This distinct integration may account for the elevated melanoma penetrance observed in this stock, with direct implications for the many laboratories employing this widely distributed model.

**SIGNIFICANCE:** This study identifies a previously undescribed Tyr::CreER^T2^ transgene integration on Chromosome 1 in JAX strain 013590, a line widely used across the melanoma research community. Accurate characterisation of this allele has direct practical implications for the many laboratories that rely on this model for studies of melanoma initiation and progression.

---

Dear Editor,

In the mouse, melanoblasts are specified from neural crest cells via SOX10-positive progenitors that delaminate from the neural tube around embryonic day 9 (E9) and subsequently upregulate expression of the master transcription factor *Mitf* followed by melanocyte lineage-restricted markers: *Pmel* and *Dct* around E10-E11, *Tyrp1* shortly thereafter and finally *Tyr* between E14.5 and E15.5 just before melanin production commences (Baxter & Pavan, 2003; Hou et al., 2000; Mackenzie et al., 1997).

The Tyr::CreER^T2^ transgenic mouse lines are critical tools for conditional gene manipulation in melanocytes, enabling tamoxifen-inducible recombination of floxed alleles specifically in pigment-producing cells. Two independent Tyr::CreER^T2^ transgenic lines are widely used in melanoma research: the Bosenberg line (Tyr::CreER^T2-Bos^; JAX strains 012328 and 013590), originally described by Bosenberg et al. (2006) and the Larue line (Tyr::CreER^T2-Lar^; JAX strain 031281), described by Yajima et al. (2006). Both use tyrosinase regulatory elements to drive melanocyte-specific CreER^T2^ but differ in construct design. The Bosenberg line combines a separated distal enhancer and proximal promoter plus a *β-globin* intron. The Larue line uses a single contiguous promoter/enhancer fragment without an intron. These lines have been widely adopted for studies of melanoma initiation and progression, including the commonly distributed Tyr::CreER^T2-Bos^;Braf^CA^;Pten^flox^ model (JAX strain 013590). JAX report that this strain exhibits higher spontaneous melanoma penetrance than originally described (*JAX - Strain 013590*, 2026).

Short-read whole genome sequencing mapped the Bosenberg transgene to a single Chromosome 2 locus, and the Larue transgene to a complex integration that generated a fusion chromosome joining proximal Chromosome 2 to distal Chromosome 7 (Aktary et al., 2020), corroborating an independent targeted locus amplification (TLA) mapping of the same Bosenberg line to the identical Chromosome 2 locus (Cain-Hom et al., 2017).

We performed long-read sequencing of a 013590 founder predicted to be homozygous for the Tyr::CreER^T2-Bos^ transgene (JAX labs qPCR) and confirmed wildtype at the *Braf* locus and homozygous for *Pten*^*flox*^ (Transnetyx), alongside a Tyr::CreER^T2^ negative control animal. First, to confirm *Pten*^*flox*^ status, Nanopore reads containing sequences flanked by a consensus loxP site (ATAACTTCGTATAGCATACATTATACGAAGTTAT) were determined using grep, a regular expression search algorithm (Thompson, 1968). A total of 7 reads matched the pattern in either the forward or the reverse direction. The sequences flanked by loxP sites were compared to the *M. musculus* genome (GRCm39/mm39, UCSC) where they aligned perfectly to chr19:32,776,614-32,778,405 - as expected the loxP sites flanked *Pten* exon five (Fig. S2).

Junction PCR at the three previously reported integration sites (method from Aktary et al., 2020) gave no transgene-specific product in any founder (Fig. S1). Reads aligning to the Bosenberg construct - including its diagnostic separated *Tyrosinase* enhancer/promoter and *β-globin* intron - were present in the dataset, establishing that this founder carries the Bosenberg rather than the Larue transgene (Fig. S3). Consistent with this, reads mapped continuously across the wild-type junction PCR sites described in Aktary et al. (2020) at the Bosenberg Chromosome 2 locus and at both breakpoints of the Larue fusion chromosome (Fig. S4-S6), with no evidence of a breakpoint at any of them. To identify its unknown integration site, we identified reads that aligned to non-mouse portions of the predicted Tyr::CreER^T2-Bos^ transgene and assembled them de novo using Flye (Kolmogorov et al., 2019), yielding multiple contigs. No contig spanned the full insertion and the longest read (63,275 bp) contained three transgene repeats. To determine precise insertion break points, we performed BLAST analysis of the above mapped reads against the *M. musculus* genome (GRCm39/mm39, UCSC). From the BLAST results, we identified two long reads that aligned to both the transgene and the genome: one spanning chr1:86,999,282–87,018,373 and one spanning chr1:87,020,805–87,034,139, suggesting that the transgene had inserted between chr1:87,018,374 and chr1:87,020,804, replacing a 2,431 bp genomic sequence. We extracted the two 1,500 bp flanking sequences (±750 bp) around the break points and used BLAST to further identify 17 reads that contained junction sequences. These reads were truncated to eliminate internal transgene repeats and *de novo* assembled using Flye (Kolmogorov et al., 2019), generating a single 63,404 bp contig with both termini mapping to Chromosome 1, confirming the integration site and 2.4 kb genomic deletion.

This region is flanked by genes *Alppl2* and *Alpi* while the deletion itself overlaps with one predicted lncRNA (*Gm29371)* of unknown function (Fig. 1). Nanopore read coverage confirmed the deletion, with wild-type controls showing continuous alignment across chr1:87,018,374-87,020,804 while homozygous transgenic animals exhibited complete absence of coverage at this locus (Fig. S7). PCR genotyping with primers spanning the 5’ deletion breakpoints (Table S1) confirmed homozygosity for the transgene insertion and that F1 animals carried one intact copy of the chromosome 1 region (Fig. S8). Genotyping of het × het progeny by qPCR (Transnetyx; Table S2) confirmed that offspring were born at Mendelian ratios (Table S3). Together, these findings indicate that the line 013590 founders provided by JAX carry a distinct Tyr::CreER^T2^ allele which may account for the observed increase in melanoma penetrance.

**Figure 1.**
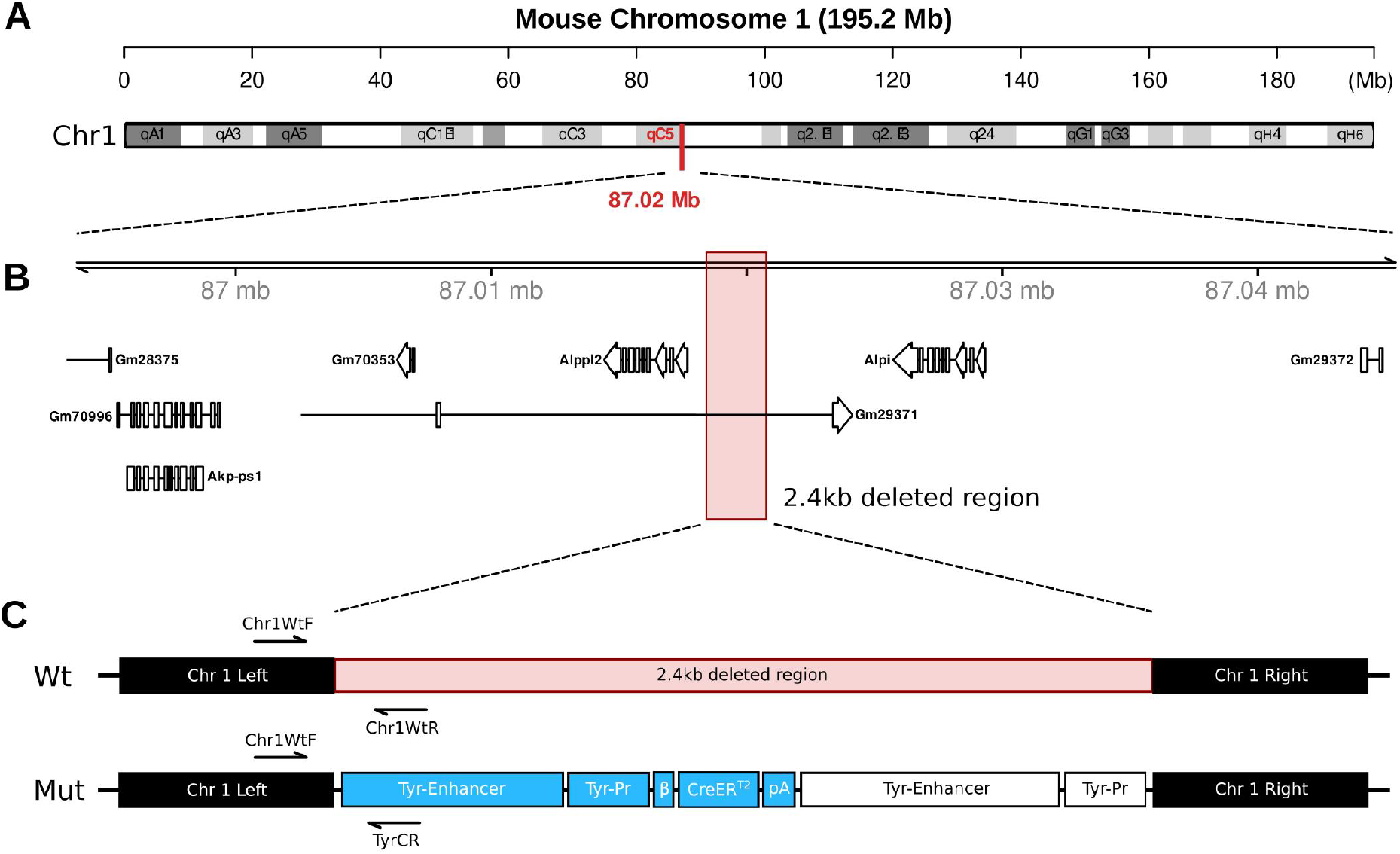
A distinct Tyr::CreER^T2^ integration of unknown copy number on Chromosome 1 in JAX strain 013590. **(A)** Whole-chromosome view of mouse Chromosome 1 (GRCm39/mm39) showing the unexpected transgene integration site at 87.02 Mb within cytogenetic band 1qC5 (red marker), distinct from the previously reported Chromosome 2 integration. **(B)** Genomic context at the integration locus reveals that the transgene replaces a 2.4 kb intergenic region (chr1:87,018,374-87,020,804, pink highlight) between *Alppl2* and *Alpi*, without disrupting coding sequences. Gene models indicate transcription direction. **(C)** Schematic of the Tyr::CreER^T2^ integration structure on wild-type (Wt) and mutant (Mut) chromosome 1. One complete transgene cassette (blue) is shown, flanked by partial transgene sequences at the 3’ end; both ends and their orientation were confirmed by nanopore sequencing spanning each junction. However, no single read spanned the entire integration locus, so the number of tandem copies is unknown - at least three are present - but only one complete copy is shown for clarity. Breakpoint-spanning primers used for junction PCR genotyping (Table S1) are indicated. Abbreviations: Tyr-Pr = *Tyr* promoter; β = *β-globin* intron; Mb = megabase, bp = base pairs; kb = kilobase; pA = poly-A signal.

The original Bosenberg founder colony comprised three retained transgenic lines (Bosenberg et al., 2006), of which only one has been molecularly characterised, by whole genome sequencing (Aktary et al., 2020) and targeted locus amplification (Cain-Hom et al., 2017), both mapping to the same Chromosome 2 locus. Neither study reported an integration on Chromosome 1, and our animals, derived from JAX strain 013590, show no evidence of the Chromosome 2 integration by junction PCR or by continuous nanopore read coverage across that site (Fig. S1, S4). The most parsimonious interpretation is that separately maintained colonies of this line carry different integrations, either because strain 013590 derives from a founder distinct from the one previously characterised, or because an original founder carried more than one integration which subsequently segregated during independent colony maintenance. We are unable to distinguish between these possibilities with the present data.

The number of tandem transgene copies at the Chromosome 1 site is also unresolved, since no single read spans the full integration locus. The integration site and the copy number may each contribute to the reported difference in penetrance and we cannot separate them. We recommend laboratories confirm which integration is present in their own stock by position-specific genotyping (e.g. junction PCR) rather than assume the previously published site. This is particularly important for melanoma studies, which often involve several additional genetic modifications and therefore complex crosses in which correct allele identification is essential.

## ACKNOWLEDGEMENTS

The authors are grateful to Prof. Lionel Larue and Dr. Zackie Aktary (Institut Curie) for predicted sequences for Tyr::CreER^T2^ transgenes. And to IT services at Lancaster University for access to The High End Computing (HEC) service. This work was supported by funding from the Medical Research Council (MR/Z504336/1) to RLM and BS.

## DATA AVAILABILITY STATEMENT

The data that support the findings of this study will be available after publication at SRA - https://www.ncbi.nlm.nih.gov/sra/ and from the corresponding author on reasonable request.

## AUTHOR CONTRIBUTIONS STATEMENT

Conceptualisation: RLM. Formal Analysis: ELW, LP, BS. Funding Acquisition: RLM. Investigation: ELW, LP, BS, AN; Methodology: RLM, BS; Project Administration: RLM; Resources: RLM; Software: BS; Supervision: RLM; Validation: LP, BS, AN; Visualisation: LP, BS, RLM, AN; Writing - Original Draft Preparation: RLM, LP, BS; Writing - Review and Editing: LP, BS, RLM, AN.

## Supplementary Methods

### Animal work

All animal work was approved by Lancaster University’s animal welfare and ethical review body (AWERB) and was performed in accordance with institutional guidelines under licence by the UK Home Office (PPL PP0041551). Mice were maintained in the animal facilities of Lancaster University. Tyr::CreER^T2-Bos^; *Pten*^*flox*^ and *Braf*^*CA*^*;Pten*^*flox*^ mice were imported from JAX Labs (strain 013590) and were maintained on a C57BL/6 background. Mice were genotyped using PCR (or qPCR by Transnetyx) as detailed below, they were housed in a barrier facility with 12-hour light and dark cycles.

### Nanopore Sequencing

Spleen tissue was dissected and stored at 4°C in Monarch StabiLyse DNA/RNA Buffer. Subsequently, high molecular weight genomic DNA was extracted using the Promega Wizard HMW DNA Extraction Kit. Long-read whole genome sequencing was performed using Oxford Nanopore’s Ligation Sequencing DNA V14 protocol (SQK-LSK114) on a Promethion flow cell (FLO-PRO114M) resulting in 12-15X genome coverage. Sequencing was conducted on the Promethion-24 instrument at Source Genomics, Cambridge, UK.

### Genotyping

gDNA was extracted from ear clip tissue in 25 mM NaOH at 98°C with gentle agitation for 1-hour followed by pH adjustment with 40 mM Tris HCl to pH 8.3. Target amplicons were amplified using Phusion Hot Start Flex DNA Polymerase (NEB, M0535S) with GC buffer and the addition of DMSO (NEB, B0515A). PCR conditions were 30 cycles: denaturation - 98 °C for 15 secs; annealing - 62 °C for 30 secs; extension - 72 °C for 15 secs. Amplified DNA was then purified by silica spin column, as per manufacturer guidance (Thermo Scientific, 10400450). Gel electrophoresis was performed using 1.5% Low EEO agarose gels (Melford 9012-36-6) in TAE buffer (40 mM Tris base, 20 mM Acetic acid, 1 mM EDTA pH 8.3) supplemented with 1X SybrSAFE (Invitrogen™, 10328162), alongside a known kDa ladder (NEB, N3238S). Gels were visualised using a BioRad Chemidoc system.

### Bioinformatics

All bioinformatics analyses were performed using default settings unless stated otherwise. Reads containing loxP consensus sequences (ATAACTTCGTATAGCATACATTATACGAAGTTAT) were identified using grep and aligned to mm39 to confirm the presence of a floxed Pten exon 5. Nanopore reads were mapped to the predicted Tyr::CreER^T2-Bos^ transgene sequence using minimap2 (v2.30; -ax map-ont). Reads mapping to the transgene (n=287) and breakpoint-spanning reads (n=17; ≥95% query coverage and >95% identity to junction sequences) were assembled using Flye (v2.9.6; --keep-haplotypes --nano-hq). Transgene-to-genome breakpoints were identified by BLAST (v2.17.0) against the mm39 reference genome (refseq_genome; accessed 11/01/2026). Junction reads were truncated to ±22,000 bp around each breakpoint prior to assembly. Read alignments were visualised in IGV (v2.19.7). Wild-type amplicon coordinates in GRCm39/mm39 were determined by in-silico PCR (UCSC) using the primer sequences of Aktary et al. (2020) (Table S1); coordinates were not lifted over from mm10.

### AI Statement

Claude (Anthropic) was used as a writing assistance tool to support text editing during manuscript preparation. It did not generate ideas, data, analyses, or conclusions.

---

**Table S1.**
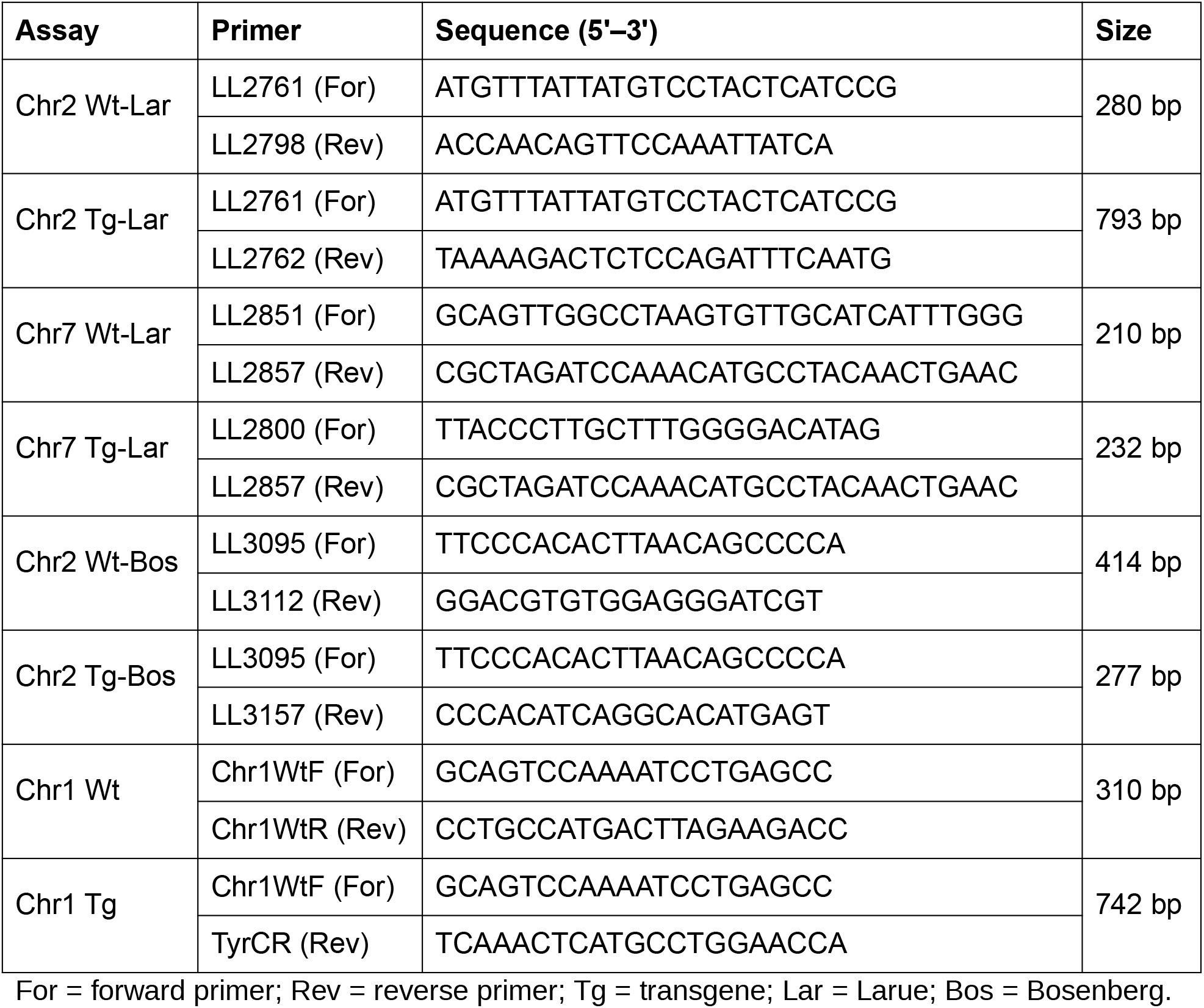
Oligonucleotide sequences used for genotyping.

**Table S2:**
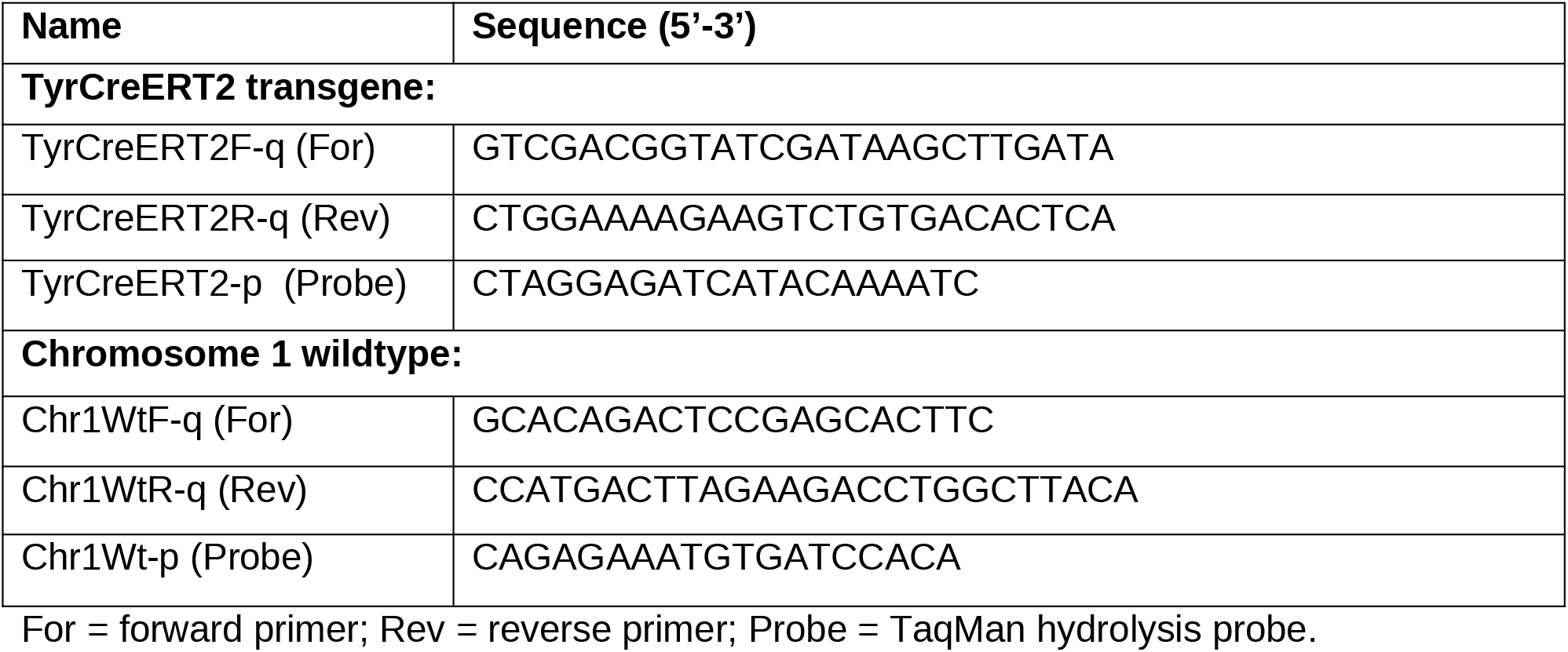
Oligonucleotide Sequences used for TaqMan qPCR Genotyping of Chromosome 1 Tyr::CreER^T2^ integration.

**Table S3:** Genotyping analysis at Chromosome 1 Tyr::CreER^T2^ integration site.

| Parent | Genotype | Observed | Expected |
| --- | --- | --- | --- |
| Het x Het | +/+ | 10 | 9.5 |
| Het x Het | +/- | 19 | 19.0 |
| Het x Het | -/- | 9 | 9.5 |
| Chi-squared | *Chi-squared = 0.05, df = 2, P = 0.98* |  |  |

Chi-squared goodness-of-fit test against expected 1:2:1 Mendelian ratio. +/+, +/-, -/- denote genotype at the Chromosome 1 Tyr::CreER<sup>T2</sup> locus.

**Supplemental Figure 1.**
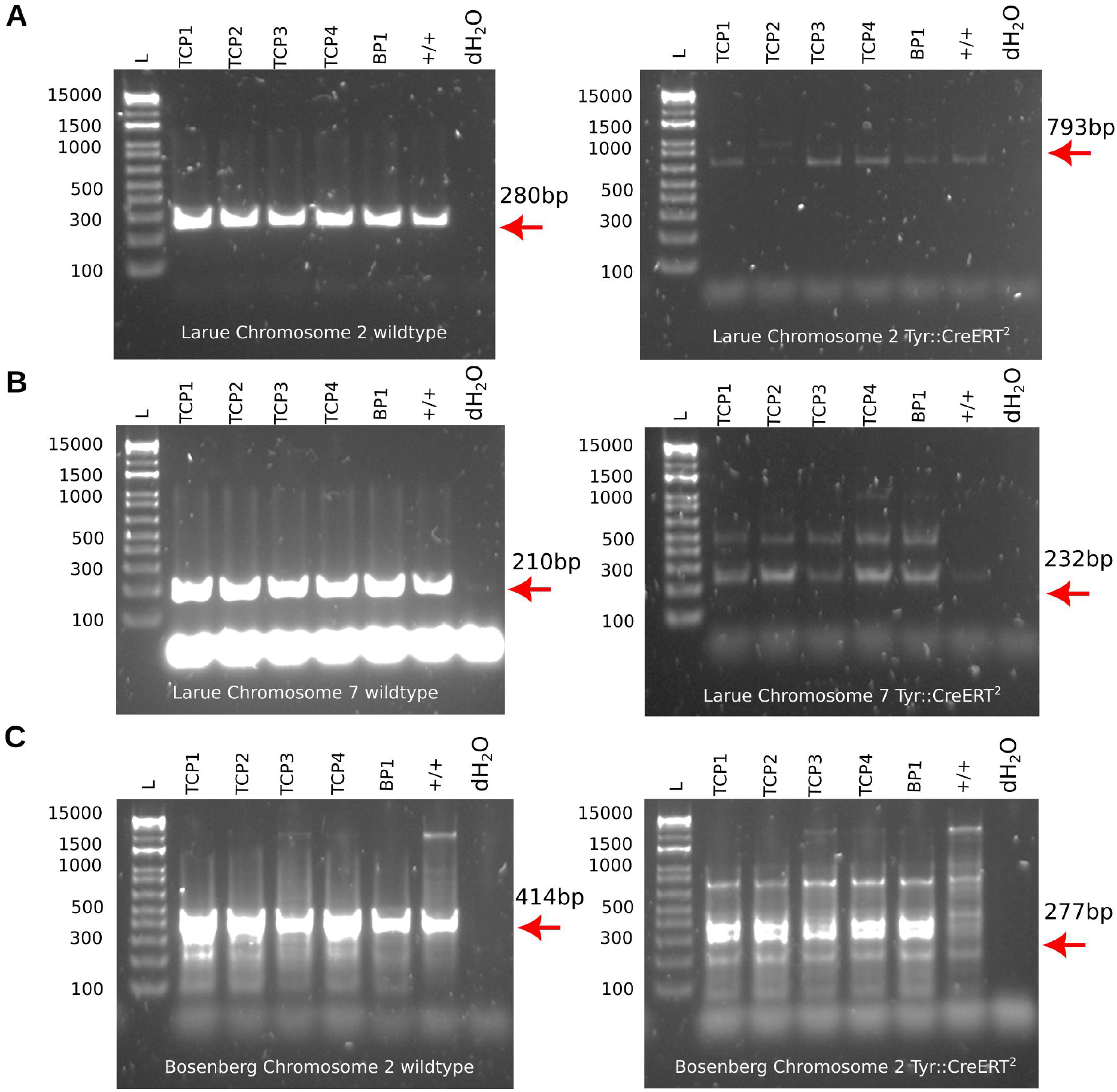
PCR genotyping of founder animals at the three previously reported Tyr::CreER^T2^ integration sites. Wild-type-specific (left) and transgene-specific (right) PCR for (A) Larue Chromosome 2, (B) Larue Chromosome 7, and (C) Bosenberg Chromosome 2 (primer sequences in Table S1). All founders (TCP1-4) and negative controls (BP1 and +/+) produced the expected wild-type band at each locus (280 bp, 210 bp, and 414 bp, respectively). No band of the correct size for the transgene-specific product (793 bp, 232 bp, and 277 bp) was observed in any sample; in (C), the band nearest the expected 277 bp product ran above 300 bp. These non-specific bands were also present in the negative controls (BP1 and +/+). Abbreviations: L, DNA ladder (bp); dH_2_O, no-template control; TCP = Tyr::CreER^T2^;*Pten*^*flox*^; BP = *Braf*^*CA*^*;Pten*^*flox*^; +/+ = wildtype control.

**Supplemental Figure 2.**
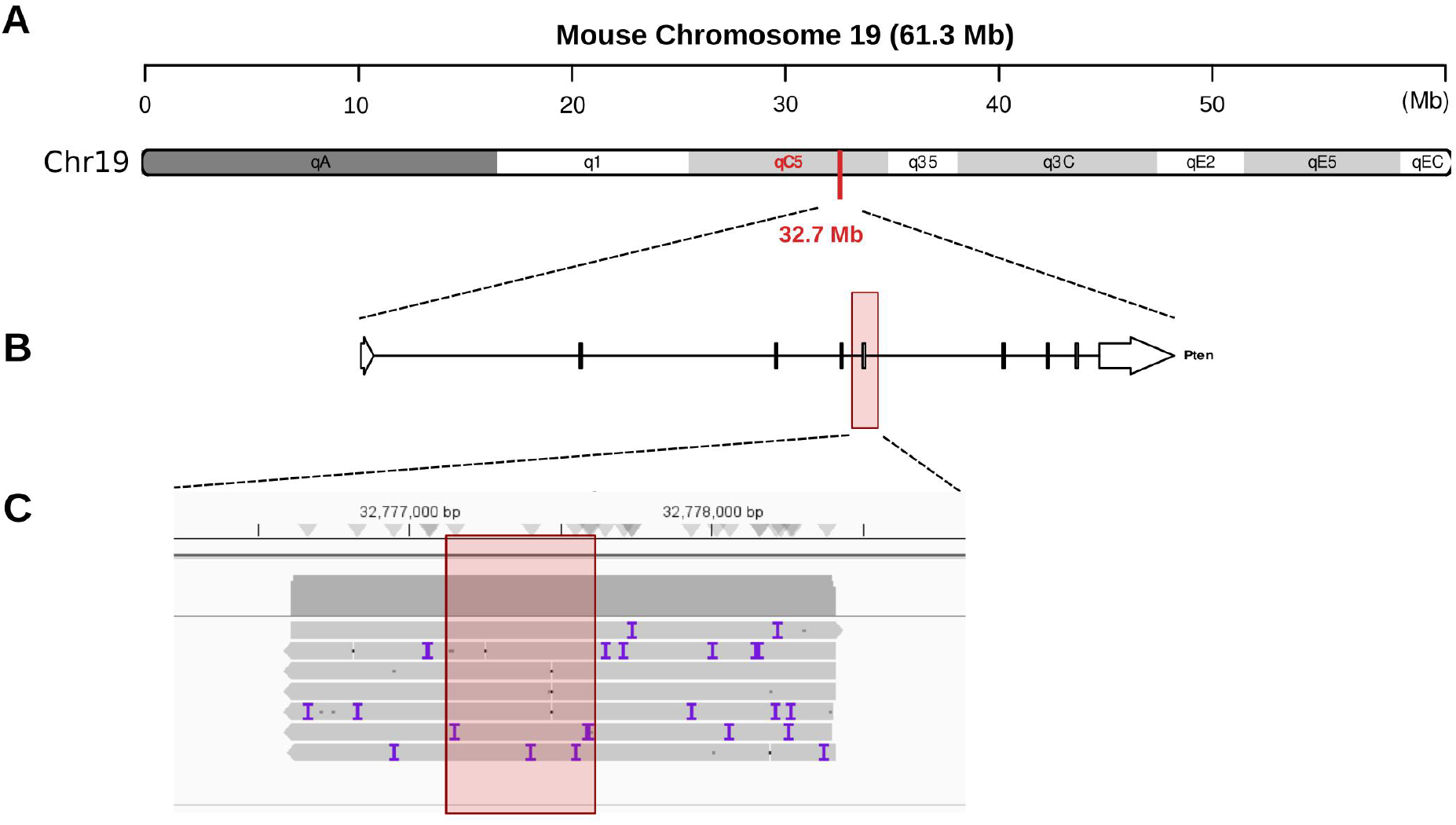
Nanopore sequencing coverage confirms the presence of a loxP-flanked *Pten* exon 5 allele. **(A)** Expanded view of chromosome 19. **(B)** Exon annotation for *Pten* **(C)** Integrative Genomics Viewer (IGV) view showing genome alignment of sequences flanked by two loxP sites, extracted from 7 reads (each indicated by a light grey horizontal bar). These loxP-flanked sequences aligned to chr19:32,776,614-32,778,405 (GRCm39/mm39), a region which encompasses *Pten* exon 5 (chr19:32,777,261-32,777,499; indicated by red box). Abbreviations: Mb = megabase.

**Supplemental Figure 3.**
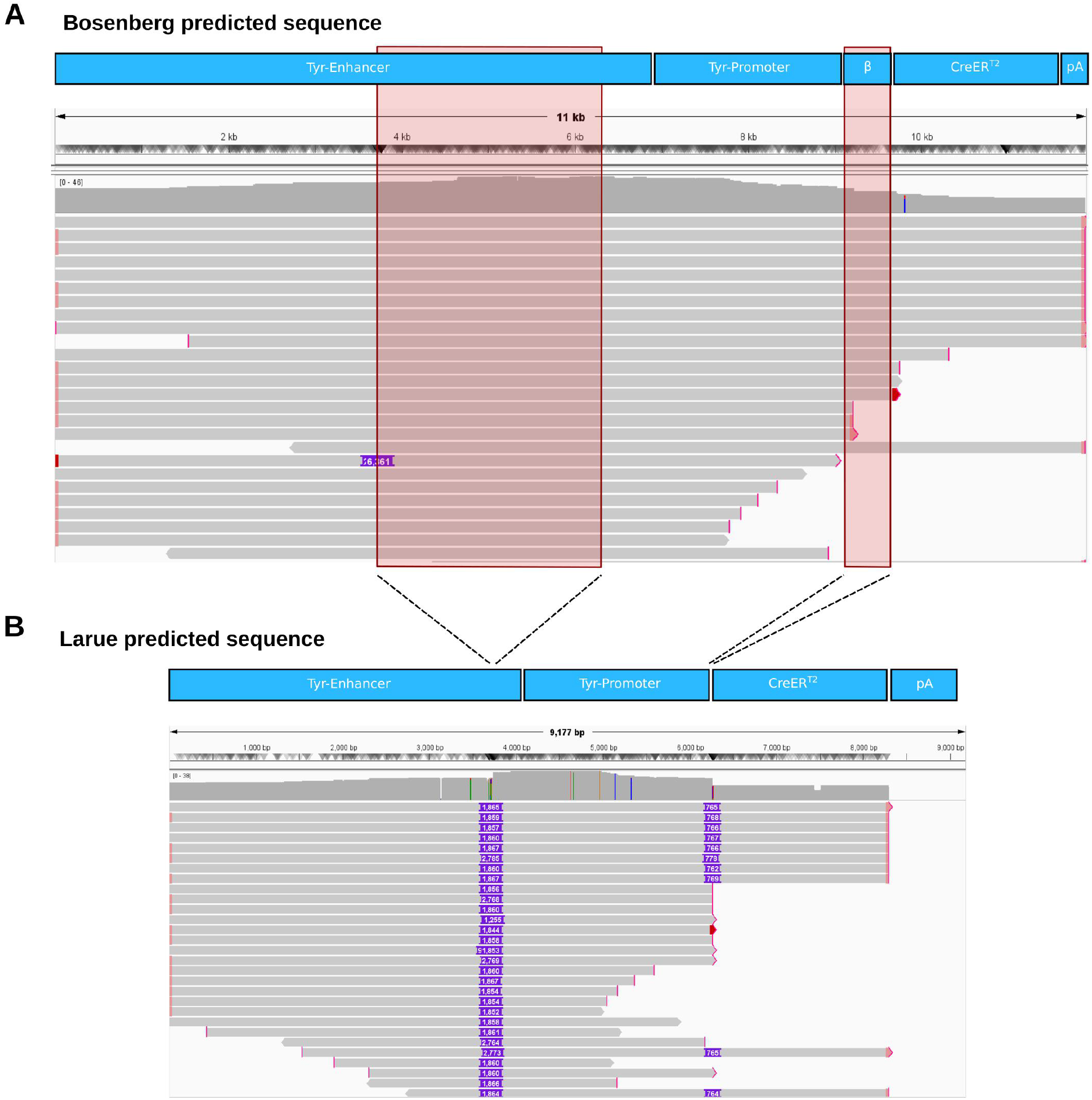
Nanopore reads confirm the presence of the Bosenberg transgene. Alignment of reads spanning the Chromosome 1 insertion against the predicted Tyr::CreER^T2-Bos^ (A) and Tyr::CreER^T2-Lar^ (B) construct sequences. Continuous read alignment covers the Bosenberg construct, including the separated *Tyr* enhancer and promoter elements and the *β-globin* intron (A). In the Larue alignment (B), gaps (red boxes in A and purple regions in B) correspond to the truncated *Tyrosinase* promoter and the absent *β-globin* intron. Abbreviations: kb = kilobase; bp = base pairs; pA = poly-A signal.

**Supplemental Figure 4.**
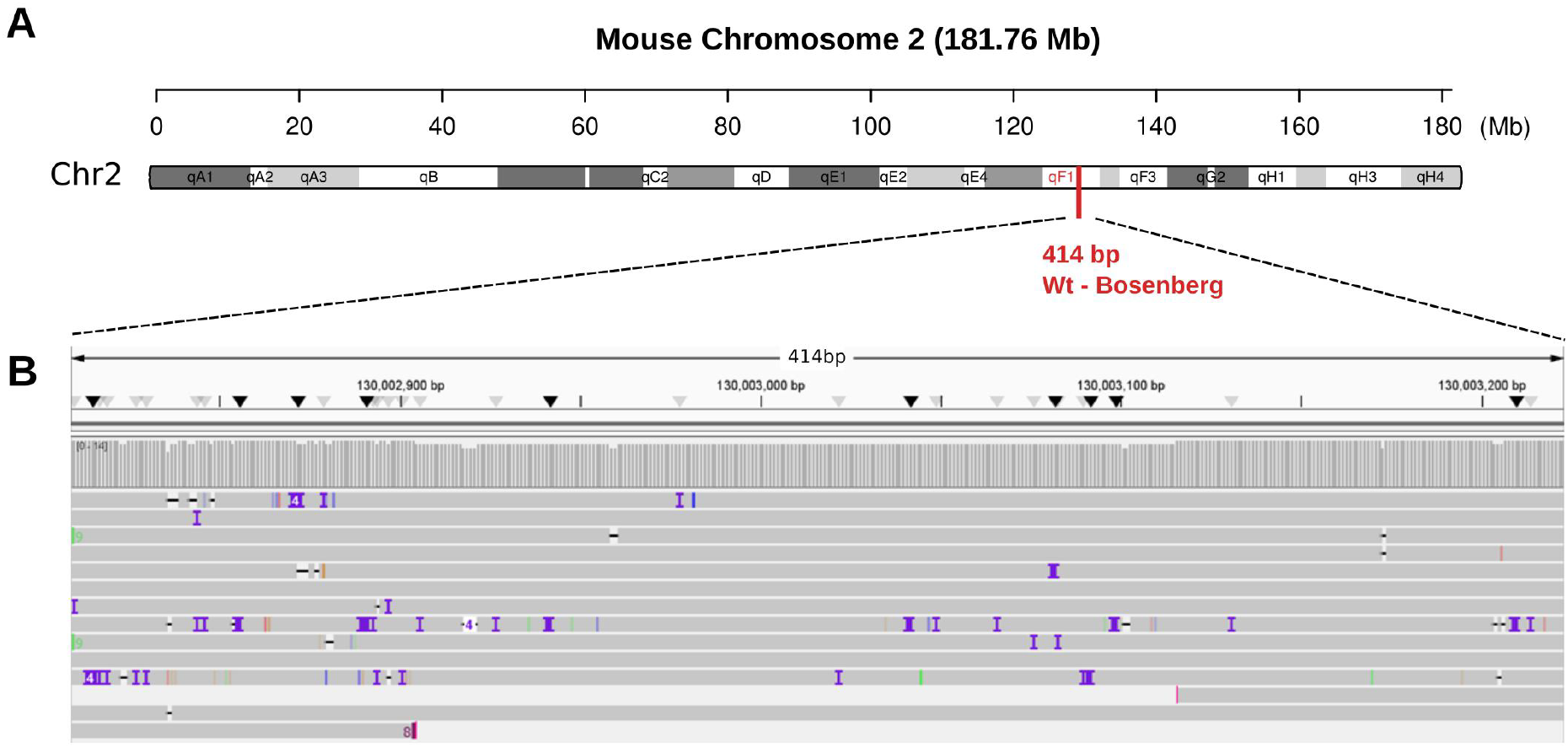
Nanopore sequencing confirms wild-type sequence at the Bosenberg Chromosome 2 wild-type junction PCR site. (A) Location of the wild-type junction PCR region (Aktary et al., 2020) on chromosome 2, cytogenetic band qF1. The wild-type amplicon spans the reported 5′ breakpoint (chr2:130,160,936, mm10), as LL3112 lies within the ∼3 kb deleted interval. (B) IGV view (540 bp shown) of read alignment across the wild-type junction PCR amplicon (chr2:130,002,809-130,003,222, 414 bp). No breakpoint was identified, consistent with absence of the Tyr::CreER^T2-Bos^ transgene at this locus. Abbreviations: Mb = megabase, bp = base pairs.

**Supplemental Figure 5.**
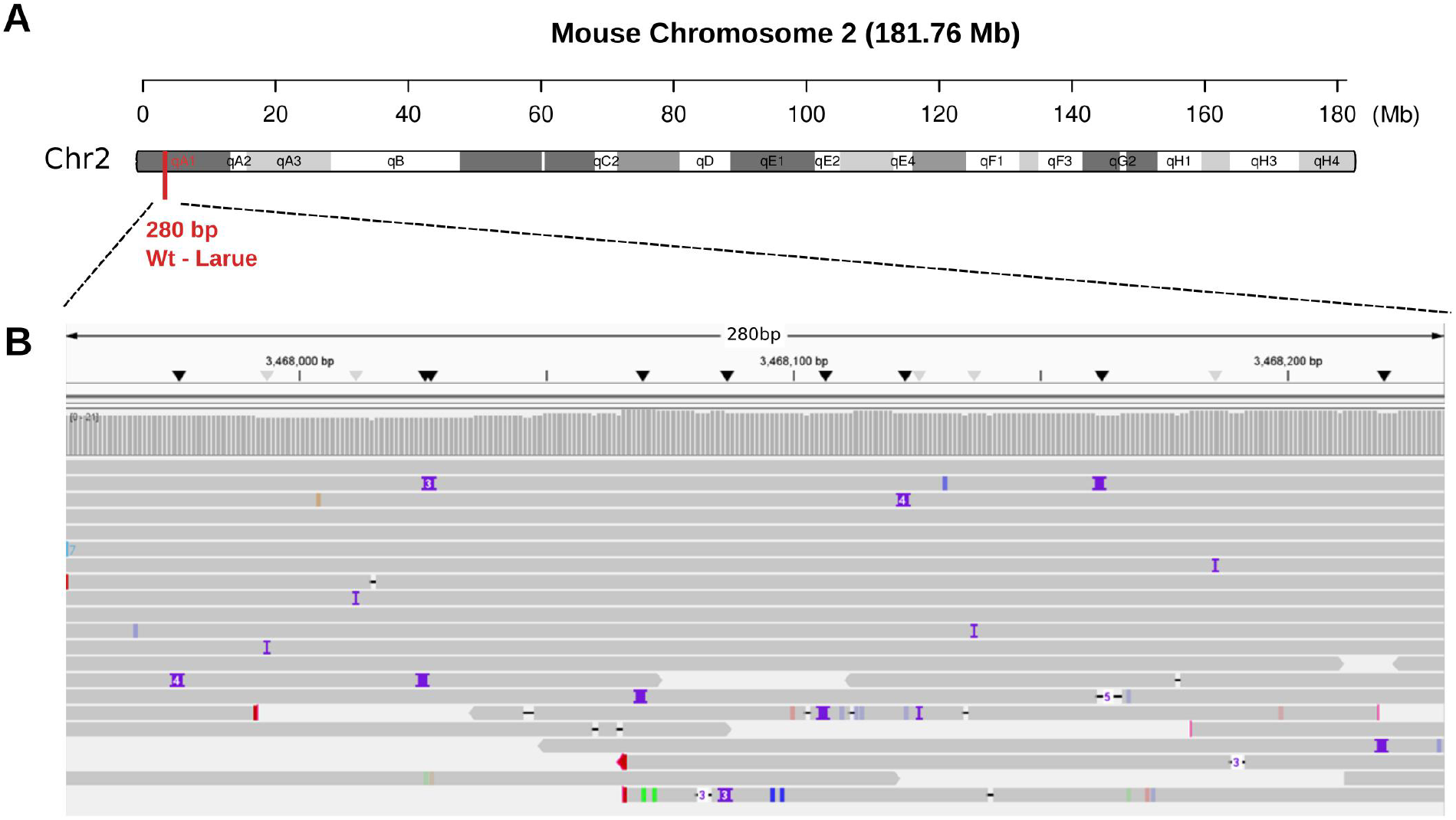
Nanopore sequencing confirms wild-type sequence at the Larue Chromosome 2 wild-type junction PCR site. (A) Location of the wild-type junction PCR region (Aktary et al., 2020) on chromosome 2, cytogenetic band 2qA1. The wild-type amplicon spans the reported breakpoint, as LL2798 lies distal to chr2:3,467,068 (mm10) and is absent from the fusion chromosome. (B) IGV view (280 bp shown) of read alignment across the wild-type junction PCR amplicon (chr2:3,467,953-3,468,231, 280 bp). No breakpoint was identified, consistent with absence of the Tyr::CreER^T2-Lar^ transgene at this locus. Abbreviations: Mb = megabase, bp = base pairs.

**Supplemental Figure 6.**
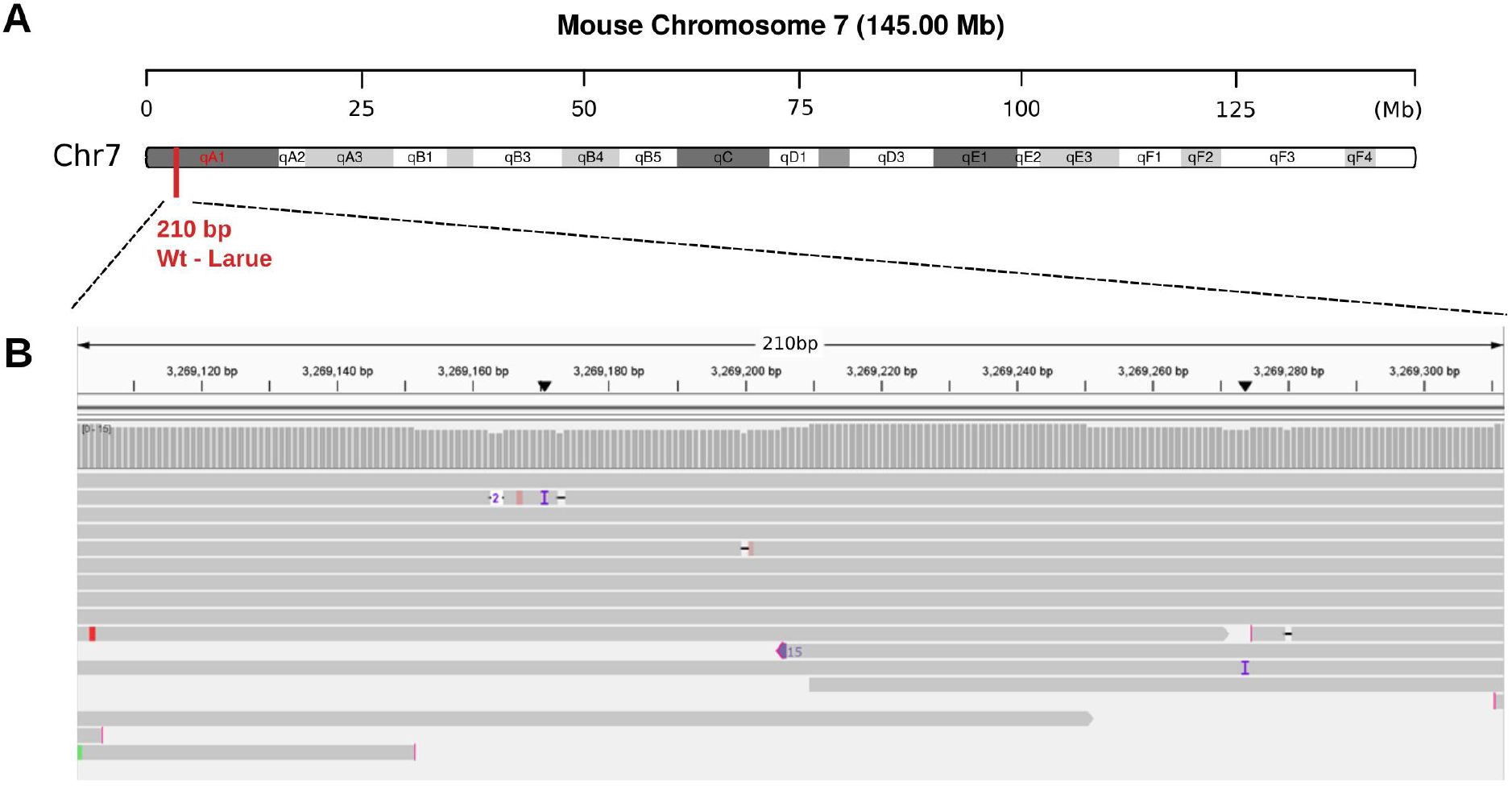
Nanopore sequencing confirms wild-type sequence at the Larue Chromosome 7 wild-type junction PCR site. (A) Location of the wild-type junction PCR region (Aktary et al., 2020) on chromosome 7, cytogenetic band 7qA1. The wild-type amplicon spans the reported breakpoint, as LL2851 lies proximal to chr7:3,220,560 (mm10) and is absent from the fusion chromosome. (B) IGV view (211 bp shown) of read alignment across the wild-type junction PCR amplicon (chr7:3,269,102-3,269,311, 210 bp). No breakpoint was identified, consistent with absence of the Tyr::CreER^T2-Lar^ transgene at this locus. Abbreviations: Mb = megabase, bp = base pairs.

**Supplemental Figure 7.**
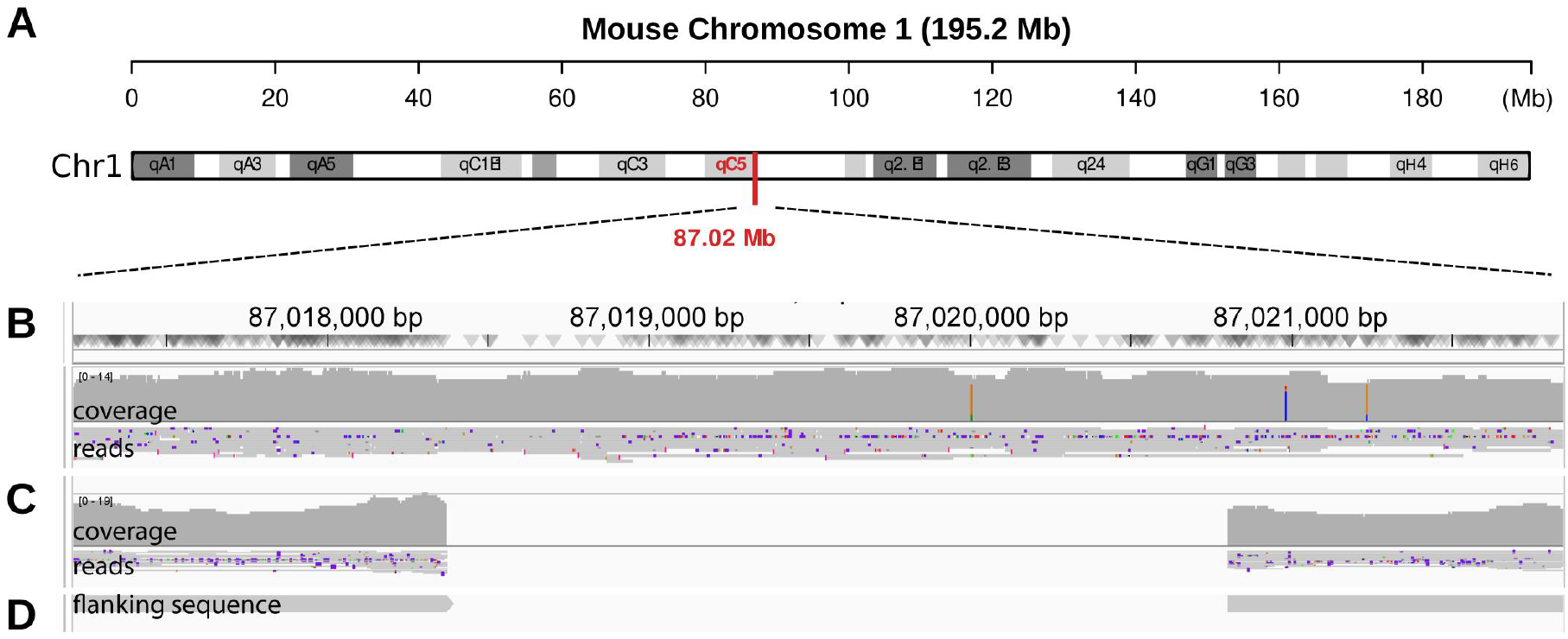
Nanopore sequencing coverage confirms the 2.4 kb deletion at the Tyr::CreER^T2^ integration site. Genome browser view of nanopore read alignments spanning chr1:87,017,209-87,021,841 (GRCm39/mm39). **(A)** Expanded view of Chromosome 1. **(B)** Wild-type control mice show continuous coverage (∼12×) across the region, including seven reads spanning the whole chr1:87,018,374-87,020,804 interval. **(C)** Homozygous Tyr::CreER^T2^ mice (JAX strain 013590) show complete absence of read coverage between the breakpoints, confirming deletion of this 2,431 bp genomic segment. Flanking regions show normal coverage with reads terminating at the integration junctions. **(D)** Consensus flanking sequences extracted from breakpoint-spanning reads, used to assemble the integration junction and design genotyping primers (Table S1). Abbreviations: Mb = megabase.

**Supplemental Figure 8.**
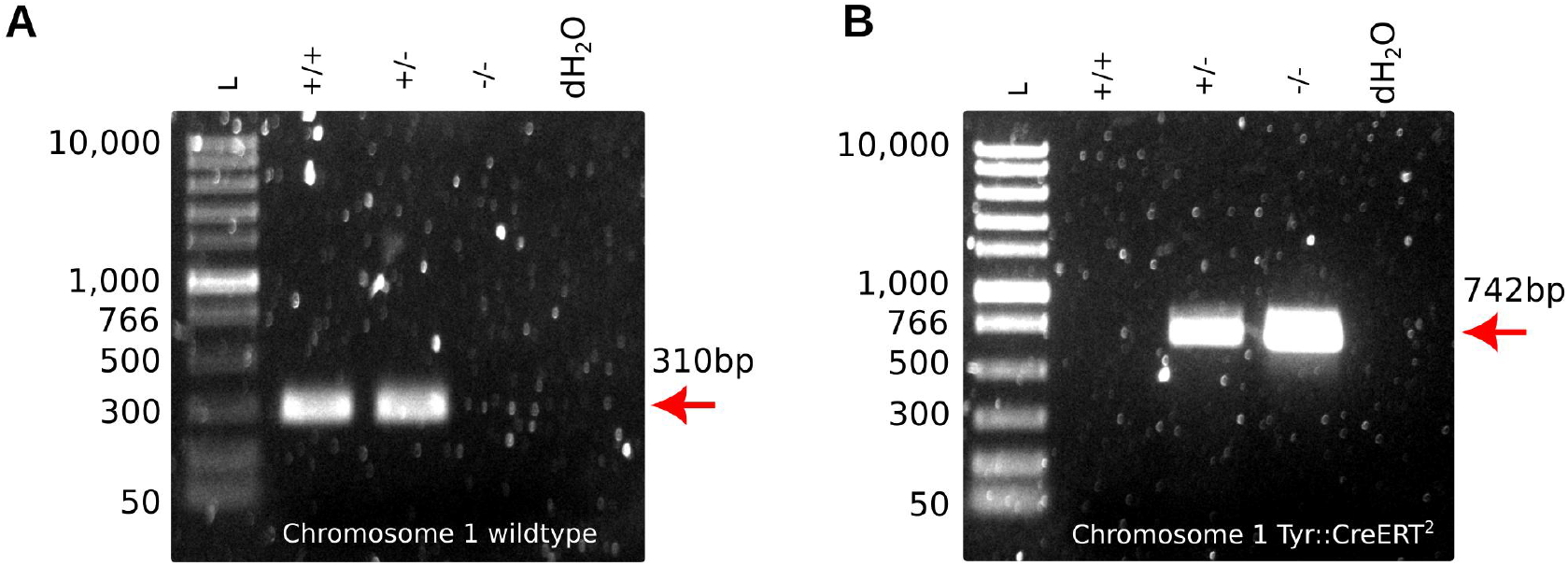
PCR genotyping assay. (A) A wildtype PCR assay using a 5’ flanking primer and a reverse primer within the 2.4 kb deletion generates a 310 bp band (Table S1, Figure 1). (B) A mutant PCR assay using a flanking primer and an internal primer with homology to the *Tyrosinase* enhancer generates a 742 bp band (Table S1, Figure 1). Band sizes are shown in base pairs. Abbreviations: L = ladder; bp = base pairs.

